# Human xenobiotic metabolism proteins have full-length and split homologs in the gut microbiome

**DOI:** 10.1101/2024.11.06.622278

**Authors:** Matthew Rendina, Peter J. Turnbaugh, Patrick H. Bradley

## Abstract

Xenobiotics, including pharmaceutical drugs, can be metabolized by both host and microbiota, in some cases by homologous enzymes. We conducted a systematic search for all human proteins with gut microbial homologs. Because gene fusion and fission can obscure homology detection, we built a pipeline to identify not only full-length homologs, but also cases where microbial homologs were split across multiple adjacent genes in the same neighborhood or operon (“split homologs”). We found that human proteins with full-length gut microbial homologs disproportionately participate in xenobiotic metabolism. While this included many different enzyme classes, short-chain and aldo-keto reductases were the most frequently detected, especially in prevalent gut microbes, while cytochrome P450 homologs were largely restricted to lower-prevalence facultative anaerobes. In contrast, human proteins with split homologs tended to play roles in central metabolism, especially of nucleobase-containing compounds. We identify twelve specific drugs that gut microbial split homologs may metabolize; two of these, 6-mercaptopurine by xanthine dehydrogenase (XDH) and 5-fluorouracil by dihydropyrimidine dehydrogenase (DPYD), have been recently confirmed in mouse models. This work provides a comprehensive map of homology between the human and gut microbial proteomes, indicates which human xenobiotic enzyme classes are most likely to be shared by gut microorganisms, and finally demonstrates that split homology may be an underappreciated explanation for microbial contributions to drug metabolism.

**Article Summary:** We develop a pipeline to systematically find human proteins with gut microbial homologs, including those split across multiple microbial genes (e.g., operons). This reveals thousands of proteins with full-length gut homologs, especially reductases and hydrolases that metabolize xenobiotics. Nearly two dozen split homologs are also observed for central metabolic enzymes, many of which can transform substrate analogs; in two cases, previous studies verify that microbial split homologs enable the expected drug to be metabolized *in vivo*. These results, which we provide as a resource, map out homology and shed light on parallel drug metabolism between host and microbiome.

## Introduction

Hundreds of small molecules, including drugs, can be metabolized by both human cells and also the trillions of microorganisms that colonize the gastrointestinal tract (the gut microbiota) [1,2]. In some cases (e.g. digoxin), drug metabolism by the microbiome can contribute to observed differences in pharmacodynamics across patients. When drugs have a narrow therapeutic window, microbial metabolism can be especially relevant, as even small differences in concentration can lead to large changes in toxicity or efficacy [3]. However, cases of microbial drug metabolism can be difficult to identify and are time- and labor- intensive to characterize; for example, metabolism of digoxin by gut microbes was first reported in 1981 [4], but the gene responsible was not identified until 2013 [5]. This has led to interest in using bioinformatic methods, such as GutBug [6], MicrobeFDT [7], and SIMMER [8], to help researchers prioritize the most promising gut microbial genes for further study.

Many proteins involved in drug metabolism are part of general “xenobiotic” systems, often with relatively broad specificity, that transform or detoxify natural products. In humans, these systems include cytochrome P450 proteins and glutathione-S-transferases. Many drugs are derived from natural products that could be encountered in the environment. These natural products typically exhibit high structural diversity, both because they are ammunition in “arms races” between competitors, and because of other constraints on natural product enzyme evolution [9]. We therefore might expect to find xenobiotic metabolic genes with less specific substrate requirements in both hosts and microbiome. In contrast, other proteins involved in drug metabolism have primary roles in central metabolism. These proteins typically have narrower specificity and metabolize drugs that are structurally similar to their natural substrates, regardless of whether they are found in nature, such as nucleoside analogs. Since many central metabolic proteins are evolutionarily ancient, one might also expect to find cases of direct homology between host and microbiome drug-metabolizing proteins. This may be especially true for drugs like chemotherapeutic or immunomodulatory antimetabolites, as these target conserved parts of metabolism.

Most approaches to detecting microbe-host homology focus on single genes. However, horizontal transfer, multidomain protein architectures, and gene fusions can complicate this picture [10–12], making it more difficult to determine whether orthologs of a human drug target or drug-metabolizing enzyme are actually likely to exist in the microbiome. Eukaryotic metabolic genes are especially likely to have bacterial origins [13], and many of these are actually fusions of bacterial operons or domains (previously termed “S-genes”). Gene fusions in eukaryotes may ensure co-expression in the absence of multi-cistronic operons, and may also function to prevent metabolic intermediates from diffusing away in a larger cell volume [12].

Taken together, this implies that gut microbial genomes may contain direct homologs to human drug metabolism genes. Some of these may be part of more general systems, while others may be central metabolic genes that happen to also metabolize designed substrate analogs. While individual cases have been identified, the full extent of such homology has remained unknown, as did whether particular systems or enzymes are particularly likely to be shared across hosts and microbes. Furthermore, in some cases, these microbial homologs may be encoded by multiple adjacent open reading frames; such “split” homologs would be missed by a one-to-one homology search. Leveraging recently-published collections of metagenome-assembled genomes (MAGs) from the human gut microbiome [14], we therefore aimed to comprehensively identify gut microbial proteins that are either full-length or “split” human homologs, then determine, based on curated human annotations, which of these were most likely to participate in xenobiotic metabolism, and which specific roles those proteins are most likely to play.

## Results

### Thousands of human proteins have either full-length or split homologs in the gut microbiome

Our approach for identifying gut microbial homologs is described in **Figure 1**. Briefly, we conducted a BLASTP homology search between gut microbial protein families and the human proteome and identified cases where a human protein aligned to ≥2/3 of a gut microbial protein. To find full-length homologs, we kept the best microbial match per genome that also aligned to ≥70% of the human protein. To find split homologs, we identified sets of gut microbial proteins that were jointly, but not individually, homologous to the majority of the human protein and encoded by adjacent or near-adjacent genes on the same strand of the same gut microbial assembly (see **Figure 1** and **Methods**).

**Figure 1:**
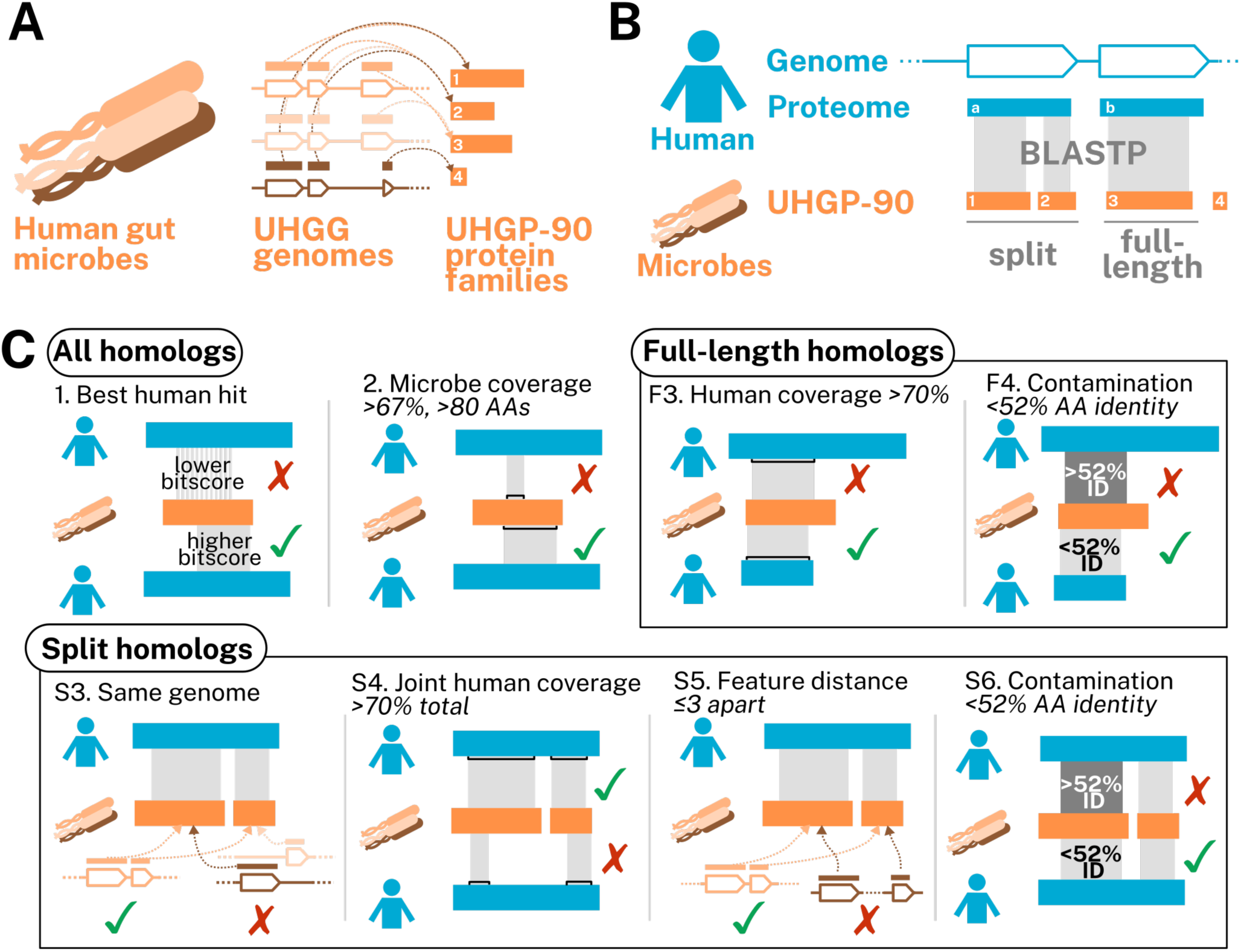
Schematic showing an overview of our approach. A) Diagram showing microbial gene and protein catalog. Human gut microbial genomes are collected in UHGG. Open reading frames (outlines, multiple shades) encode protein sequences (solid orange blocks, multiple shades). These are clustered at 90% amino acid identity (dashed lines) to form the UHGP-90 protein families (solid orange blocks, single shade). B) Pipeline overview. BLASTP is used to identify cases where microbial UHGP-90 proteins jointly or individually align to human proteins. Proteins that jointly align and are encoded by nearby features on the same genome are termed “split homologs.” C) Filtering criteria. Steps 1 and 2 involve finding human proteins (“best human hit”) that align to at least two-thirds of a microbial protein (“microbe coverage”). These steps are common to both pipelines (“all homologs”). For full-length homologs, alignments are then filtered based on whether they individually cover ≥70% of a human protein (“human coverage”) at less than 52% amino acid identity (“contamination”). For split homologs, bacterial proteins aligning to a human protein must be encoded in the same genome (“same genome”), cover ≥70% of the human protein (“joint human coverage”), be at most three features apart (“feature distance”), and have <52% amino acid identity (“contamination”). Note that in the “split homologs” section, step 3 is re-run after steps 4, 5, and 6. See **Methods** for details.

Our gut microbial sequences came from the Universal Human Gut Proteome database (UHGP) [14], which contains predicted protein sequences from >200K isolate and metagenome-assembled genomes. Because UHGP contains a very large number of non-identical protein sequences (>170M), we used a derivative of UHGP clustered at 90% amino acid identity (UHGP-90), which retains most of the sequence diversity at less than one-tenth the size (14M protein families) [15]. We then performed a local alignment search using BLASTP [16] for each UHGP-90 protein cluster. Because we wanted to compare our results against multiple databases, we did not limit our initial search to only known drug metabolism genes, but instead searched against all >20K human protein sequences from UniProt [15], and then filtered according to the criteria above.

Overall, we found that homology between the human and gut microbial proteomes was not rare, with 12.9% of human proteins (2,589) having at least one gut microbial homolog. Furthermore, while the majority had full-length homologs, a sizable minority (313) had at least one split homolog. In fact, 23 human proteins had more split than full-length homologs, and 16 had no full-length homologs at all (**Figure 2**, **Table 1**), meaning that they could not have been found by a conventional one-to-one homology search. These numbers are similar in magnitude to a previous estimate of eukaryotic gene families that likely descended from fusions of prokaryotic proteins or domains. This study identified 282 such families, 19 of which were both widely distributed in eukaryotes and also “operon-like” in bacteria, in that they appeared in an annotated operon in at least one bacterial genome [12].

**Figure 2:**
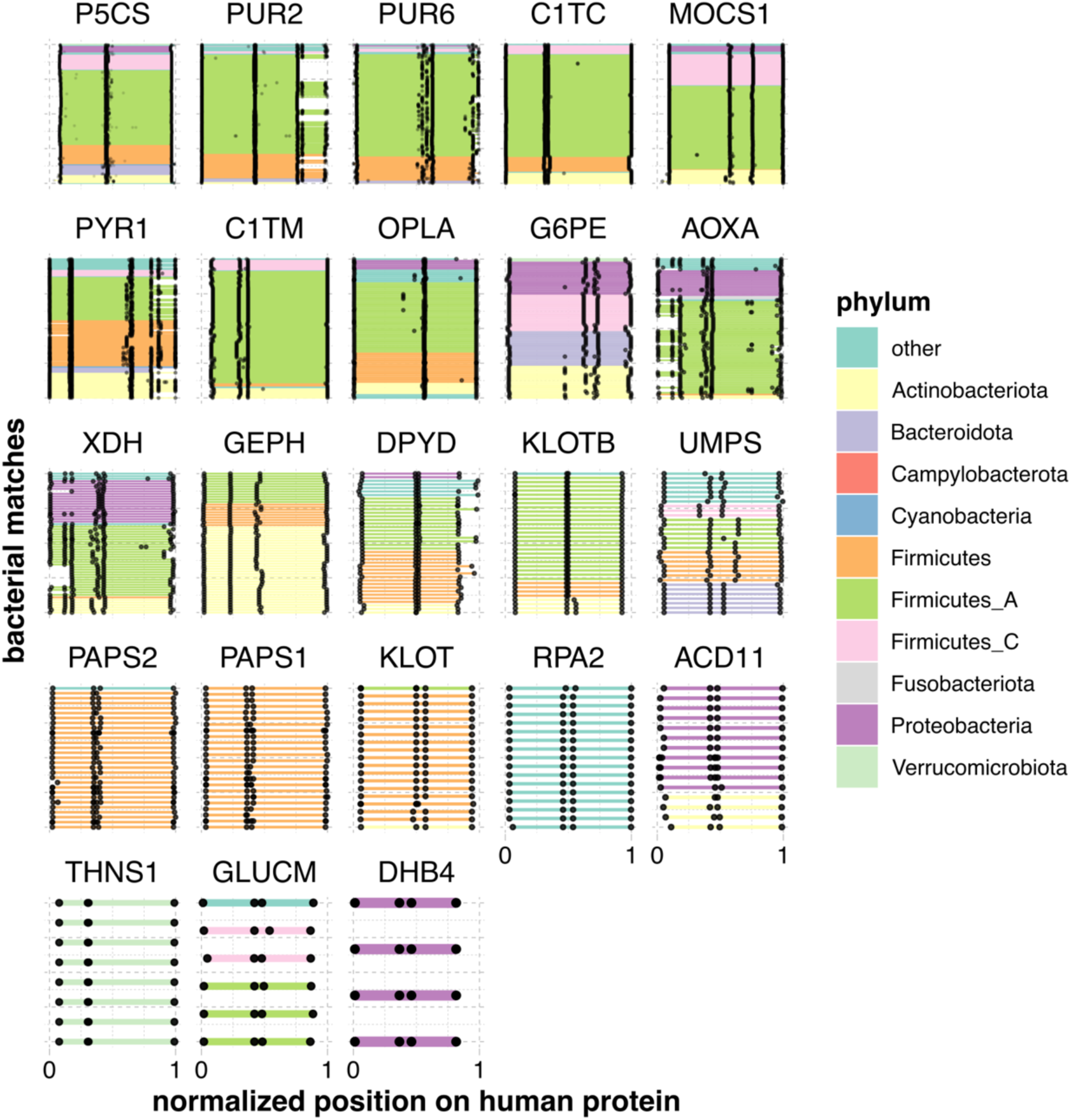
Gut microbial split homologs aligned to human proteins. Each panel shows alignments of sets of gut microbial sequences (see **Table 2**) to a single human protein. Each bacterial gene is a line segment, colored by phylum and bounded by black dots. Bacterial genes from the same genome are on the same y-axis position. Only microbial sequences that were found to be part of a neighborhood were retained, and only human proteins with more neighborhood than full-length orthologs are shown. Because both the number of microbial homologs per human protein and the lengths of the human protein sequences vary, coordinates have been transformed to between 0 and 1.

**Figure 3:**
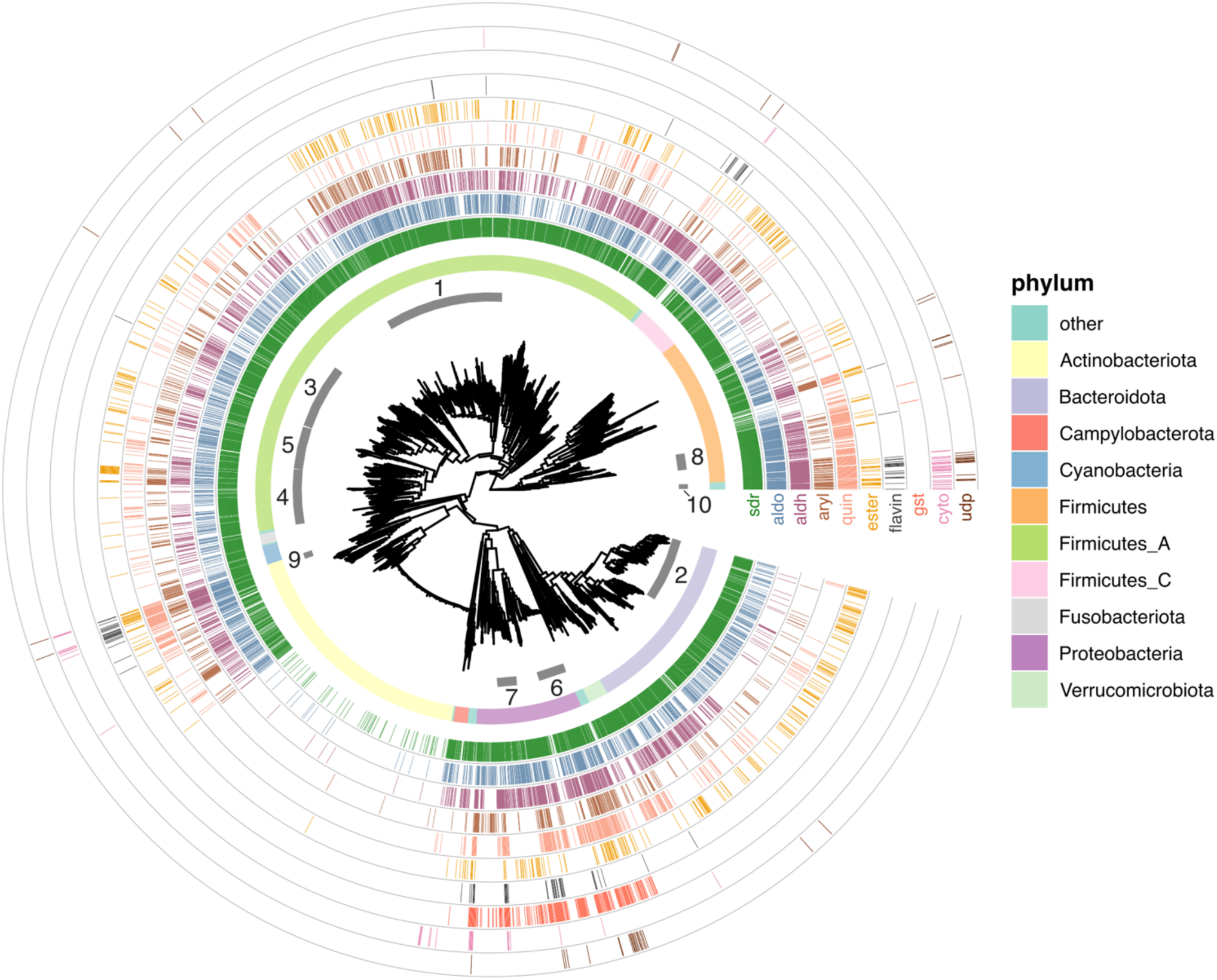
Phylogenetic distribution of full-length gut microbial xenobiotic homologs. The inset is a midpoint-rooted tree of all bacteria included in UHGG v1.0. Numbered ring segments indicate selected bacterial families (see Tables 2-3). The first complete ring shows the phylum-level classification. Note that “Firmicutes” contains Bacilli, “Firmicutes_A” contains Clostridia, and “Firmicutes_C” contains Negativicutes. Successive ring tracks mark the species where full-length homologs were identified (colored lines). From inner to outer, these are: short-chain reductases (“sdr”, dark green); aldo-keto reductases (“aldo”, blue-green); aldehyde dehydrogenases (“aldh”, purple); arylamine and arylacetamide metabolism (“aryl”, brown); quinone oxidoreductases (“quin”, salmon); type B carboxylesterases (“ester”, orange), flavin-containing mono-oxygenases (“flavin”, dark gray), glutathione-S-transferases (“gst”, red), cytochromes (“cyto”, violet), and UDP-glucuronosyl-transferases (“udp”, dark brown). Ring tracks are plotted in order of Faith’s phylogenetic diversity (PD), descending from inside to outside. Numbered families: 1. *Lachnospiraceae*, 2. *Bacteroidaceae*, 3. *Oscillospiraceae*, 4. *Acutalibacteriaceae*, 5. *Ruminococcaceae*, 6. *Enterobacteriaceae*, 7. *Burkholderiaceae*, 8. *Lactobacillaceae*, 9. *Mycobacteriaceae*, 10. *Paenibacillaceae*.

**Table 1:** Table showing all human proteins with more split than full-length homologs (nSplit, nFull) in gut bacteria.

| Entry | Entry Name | Description | nFull | nSplit |
| --- | --- | --- | --- | --- |
| P54886 | P5CS_HUMAN | Delta-1-pyrroline-5-carboxylate synthase | 27 | 293 |
| P22102 | PUR2_HUMAN | Trifunctional purine biosynthetic protein adenosine-3 | 0 | 200 |
| P22234 | PUR6_HUMAN | Bifunctional phosphoribosylaminoimidazole carboxylase/phosphoribosylaminoimidazole succinocarboxamide synthetase | 0 | 145 |
| P27708 | PYR1_HUMAN | Multifunctional protein CAD | 0 | 120 |
| Q9NZB8 | MOCS1_HUMAN | Molybdenum cofactor biosynthesis protein 1 | 0 | 75 |
| P11586 | C1TC_HUMAN | C-1-tetrahydrofolate synthase, cytoplasmic | 0 | 64 |
| O14841 | OPLA_HUMAN | 5-oxoprolinase | 7 | 54 |
| O95479 | G6PE_HUMAN | GDH/6PGL endoplasmic bifunctional protein | 7 | 47 |
| Q6UB35 | C1TM_HUMAN | Monofunctional C1-tetrahydrofolate synthase, mitochondrial | 4 | 47 |
| Q06278 | AOXA_HUMAN | Aldehyde oxidase | 3 | 46 |
| P47989 | XDH_HUMAN | Xanthine dehydrogenase/oxidase | 0 | 38 |
| O95340 | PAPS2_HUMAN | Bifunctional 3'-phosphoadenosine 5'-phosphosulfate synthase 2 | 0 | 28 |
| Q9NQX3 | GEPH_HUMAN | Gephyrin | 0 | 27 |
| Q12882 | DPYD_HUMAN | Dihydropyrimidine dehydrogenase | 24 | 26 |
| O43252 | PAPS1_HUMAN | Bifunctional 3'-phosphoadenosine 5'-phosphosulfate synthase 1 | 0 | 24 |
| P11172 | UMPS_HUMAN | Uridine 5'-monophosphate synthase | 0 | 22 |
| Q86Z14 | KLOTB_HUMAN | Beta-klotho | 0 | 12 |
| Q709F0 | ACD11_HUMAN | Acyl-CoA dehydrogenase family member 11 | 0 | 11 |
| Q7Z3D6 | GLUCM_HUMAN | D-glutamate cyclase, mitochondrial | 0 | 6 |
| Q9H9Y6 | RPA2_HUMAN | DNA-directed RNA polymerase I subunit RPA2 | 2 | 5 |
| Q9UEF7 | KLOT_HUMAN | Klotho | 0 | 5 |
| P51659 | DHB4_HUMAN | Peroxisomal multifunctional enzyme type 2 | 0 | 1 |
| Q8IYQ7 | THNS1_HUMAN | Threonine synthase-like 1 | 0 | 1 |

**Table 2:** Number of unique species per family in UHGG (rows) where at least one homolog in a particular xenobiotic enzyme class (columns) was detected. Highlighted cells show the top three families per xenobiotic class.

| phylum | family | sdr | aldo | aldh | gdxg | quin | ester | aryl | flavin | gst | cyto | udp |
| --- | --- | --- | --- | --- | --- | --- | --- | --- | --- | --- | --- | --- |
| Firmicutes_A | Lachnospiraceae | 192 | 178 | 170 | 129 | 34 | 105 | 27 | 4 | 0 | 1 | 0 |
| Bacteroidota | Bacteroidaceae | 116 | 91 | 26 | 18 | 26 | 67 | 4 | 0 | 0 | 0 | 0 |
| Proteobacteria | Enterobacteriaceae | 108 | 105 | 108 | 63 | 107 | 55 | 62 | 8 | 107 | 1 | 19 |
| Firmicutes | Lactobacillaceae | 59 | 59 | 49 | 11 | 48 | 3 | 0 | 0 | 0 | 0 | 0 |
| Firmicutes_A | Ruminococcaceae | 54 | 47 | 34 | 26 | 10 | 21 | 5 | 0 | 0 | 0 | 0 |
| Firmicutes_A | Acutalibacteraceae | 45 | 35 | 31 | 21 | 18 | 14 | 2 | 0 | 0 | 0 | 0 |
| Firmicutes_A | Oscillospiraceae | 43 | 36 | 37 | 18 | 11 | 17 | 11 | 1 | 1 | 0 | 1 |
| Firmicutes_A | Clostridiaceae | 31 | 22 | 30 | 12 | 5 | 2 | 0 | 8 | 0 | 3 | 1 |
| Campylobacterota | Campylobacteraceae | 25 | 6 | 21 | 1 | 0 | 1 | 0 | 0 | 0 | 6 | 0 |
| Proteobacteria | Burkholderiaceae | 22 | 16 | 22 | 9 | 13 | 12 | 2 | 7 | 21 | 4 | 0 |
| Firmicutes | Enterococcaceae | 20 | 20 | 18 | 12 | 20 | 5 | 3 | 1 | 1 | 0 | 0 |
| Actinobacteriota | Mycobacteriaceae | 19 | 20 | 20 | 10 | 20 | 17 | 2 | 9 | 0 | 2 | 0 |
| Firmicutes_I | Paenibacillaceae | 19 | 19 | 18 | 12 | 19 | 12 | 6 | 2 | 0 | 7 | 3 |
| Firmicutes | Staphylococcaceae | 14 | 14 | 14 | 3 | 13 | 10 | 14 | 1 | 0 | 0 | 3 |
| Proteobacteria | Moraxellaceae | 11 | 9 | 11 | 9 | 11 | 0 | 1 | 8 | 11 | 0 | 0 |
| Bacteroidota | Tannerellaceae | 11 | 11 | 10 | 2 | 7 | 5 | 2 | 0 | 0 | 0 | 4 |
| Proteobacteria | Pseudomonadaceae | 10 | 9 | 10 | 6 | 10 | 1 | 2 | 7 | 10 | 1 | 3 |
| Firmicutes | Amphibacillaceae | 7 | 7 | 7 | 0 | 7 | 1 | 3 | 1 | 0 | 6 | 0 |
| Firmicutes | Bacillaceae | 4 | 4 | 4 | 0 | 4 | 4 | 2 | 4 | 0 | 4 | 4 |
| Firmicutes | Bacillaceae_G | 4 | 4 | 4 | 3 | 4 | 0 | 4 | 4 | 0 | 3 | 4 |
| Proteobacteria | Xanthobacteraceae | 3 | 3 | 3 | 3 | 3 | 1 | 2 | 3 | 3 | 3 | 1 |

### Human proteins with full-length vs. split homologs differ in function and subcellular localization

After identifying human proteins with full-length and split homologs, we used Gene Ontology (GO) enrichment [17,18] to ask whether these two groups could be differentiated in localization and function (**Supplementary Tables 1-2**). Considering proteins with mostly full- length homologs, we observed that many of the enriched pathways were mitochondrial, such as “tricarboxylic acid cycle” (*p*_*adj*_ = 1.2 × 10^-13^), “fatty acid beta-oxidation” (*p*_*ad*j_ = 2.1 × 10^-10^), and “carnitine metabolic process” (*p*_*ad*j_ = 1.9 × 10^-16^). Because the eukaryotic mitochondrion descends from a bacterial ancestor, we might expect human proteins with gut bacterial homologs to localize to the mitochondrion. Indeed, human proteins with full-length homologs were much more likely to localize to the mitochondrion (odds ratio 4.8, 95% CI [4.3, 5.5], *p* < 2.2 × 10^-16)^, Fisher’s exact test). Further, this enrichment increased the more frequently the full-length homologs were detected (**Supplementary Figure 1**). This set of enrichments aligns strongly with previous work that identified a set of nuclear gene families present in the last eukaryotic common ancestor that had mainly Alphaproteobacterial origins, mitochondrial localization, and roles in energy production [19].

Remarkably, the most-enriched term among proteins with full-length homologs was “xenobiotic metabolism” (*p*_*ad*j_ = 8.9 × 10^-25^). There was an equally strong enrichment when considering only non-mitochondrial genes. Further, the enrichment was not driven by a single enzyme family. We observed homologs of short-chain and aldo-keto reductases, carboxylesterases, arylamine N-acetyltransferases and arylacetamide deacetylases, glutathione-S-transferases, flavin mono-oxygenases, UDP-glucuronosyl transferases and cytochrome P450 family members, among others.

We next compared these results with the proteins that had mainly split homologs. In contrast, these were not at all enriched for the “xenobiotic metabolism” GO term (*p*_*ad*j_ = 1), but rather for a smaller number of central pathways, namely purine, pyrimidine, and cofactor (folate and molybdopterin) metabolism (*p*_*ad*j_ ≤ 0.05). Full-length homologs were also significantly enriched for some of these pathways (**Supplementary Table 2**), but less so than the general “xenobiotic metabolism” term, indicating that proteins with split homologs are a more functionally specific group. Proteins in these central pathways, however, still make important contributors to drug metabolism. Nucleoside and folate analogs, in particular, are common antiviral, antibiotic, and chemotherapeutic agents.

Further, when we examined the subcellular distribution of human proteins with mainly split homologs, the fraction localizing to the mitochondria was more modest, and did not differ significantly from the base rate (odds ratio 2.0, 95% CI [0.23, 8.8], *p* = 0.29). If anything, the proteins with the most split homologs were the least likely to be mitochondrial (**Supplementary Figure 1**). Proteins with split homologs therefore appear to participate in different biological processes (cytosolic, primarily central metabolism) than full-length homologs (mitochondrial, energy production, both xenobiotic and central metabolism).

### Reductases and hydrolases dramatically outnumber cytochromes and UDP-glucuronosyltransferases in gut microbes

The above analysis indicates the presence of full-length homologs of xenobiotic metabolism enzymes in gut microbes. However, it does not tell us about their phylogenetic distribution, which is important because gut microbial clades vary in their prevalence and average abundance across orders of magnitude [20]. We therefore identified eleven enzyme families with at least some members known to participate in human xenobiotic metabolism, then determined which gut microbial species contained homologs of these families (**Figure 2**, **Table 2**), as well as how many distinct bacterial proteins were identified (**Table 3**).

**Table 3:**
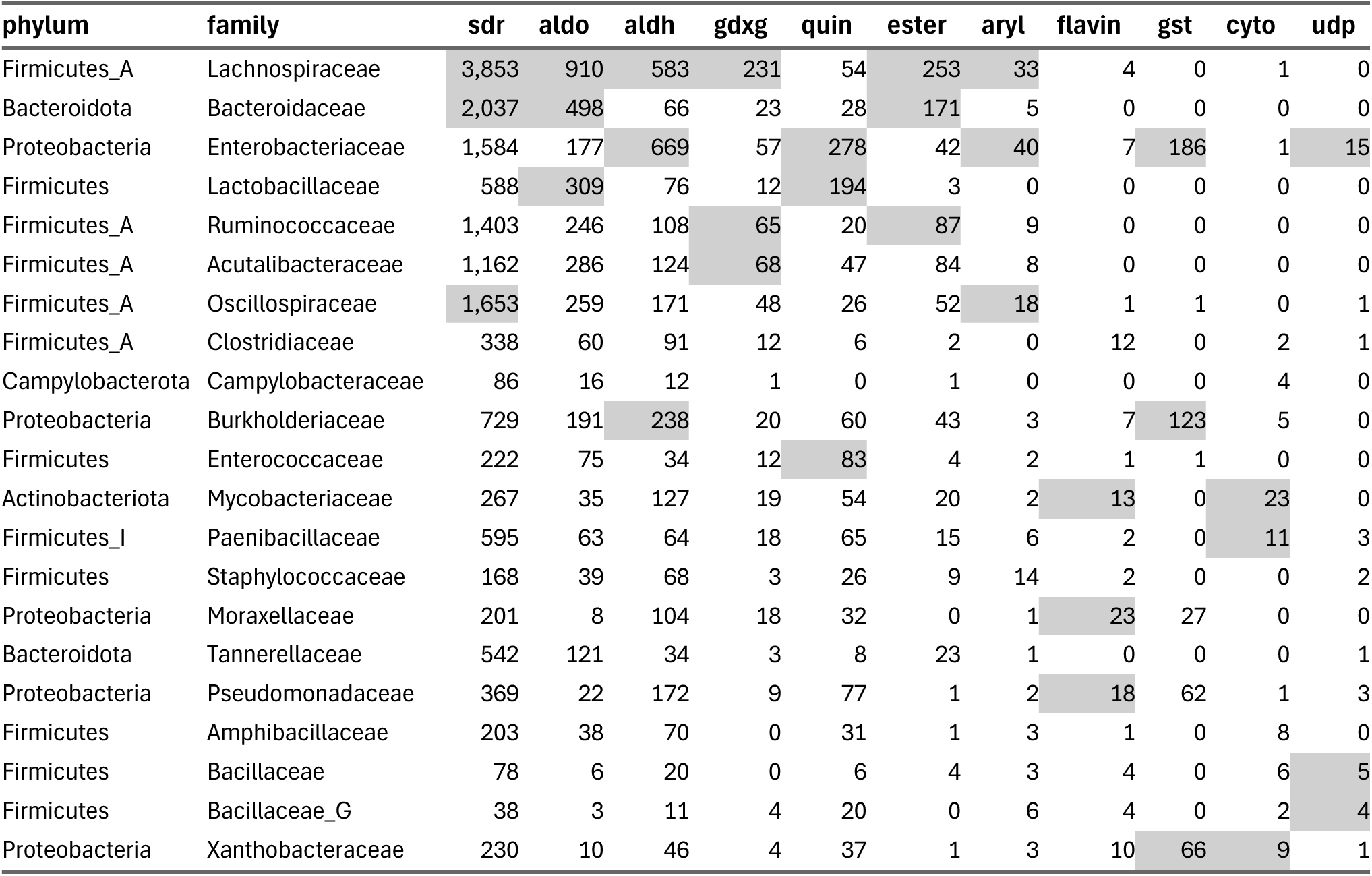
Number of unique bacterial homologs (i.e., distinct UHGP-90 protein families) of proteins in a particular xenobiotic class (columns) detected per bacterial family in UHGG (rows). Highlighted cells show the top three families per xenobiotic class.

| phylum | family | sdr | aldo | aldh | gdxg | quin | ester | aryl | flavin | gst | cyto | udp |
| --- | --- | --- | --- | --- | --- | --- | --- | --- | --- | --- | --- | --- |
| Firmicutes_A | Lachnospiraceae | 3,853 | 910 | 583 | 231 | 54 | 253 | 33 | 4 | 0 | 1 | 0 |
| Bacteroidota | Bacteroidaceae | 2,037 | 498 | 66 | 23 | 28 | 171 | 5 | 0 | 0 | 0 | 0 |
| Proteobacteria | Enterobacteriaceae | 1,584 | 177 | 669 | 57 | 278 | 42 | 40 | 7 | 186 | 1 | 15 |
| Firmicutes | Lactobacillaceae | 588 | 309 | 76 | 12 | 194 | 3 | 0 | 0 | 0 | 0 | 0 |
| Firmicutes_A | Ruminococcaceae | 1,403 | 246 | 108 | 65 | 20 | 87 | 9 | 0 | 0 | 0 | 0 |
| Firmicutes_A | Acutalibacteraceae | 1,162 | 286 | 124 | 68 | 47 | 84 | 8 | 0 | 0 | 0 | 0 |
| Firmicutes_A | Oscillospiraceae | 1,653 | 259 | 171 | 48 | 26 | 52 | 18 | 1 | 1 | 0 | 1 |
| Firmicutes_A | Clostridiaceae | 338 | 60 | 91 | 12 | 6 | 2 | 0 | 12 | 0 | 2 | 1 |
| Campylobacterota | Campylobacteraceae | 86 | 16 | 12 | 1 | 0 | 1 | 0 | 0 | 0 | 4 | 0 |
| Proteobacteria | Burkholderiaceae | 729 | 191 | 238 | 20 | 60 | 43 | 3 | 7 | 123 | 5 | 0 |
| Firmicutes | Enterococcaceae | 222 | 75 | 34 | 12 | 83 | 4 | 2 | 1 | 1 | 0 | 0 |
| Actinobacteriota | Mycobacteriaceae | 267 | 35 | 127 | 19 | 54 | 20 | 2 | 13 | 0 | 23 | 0 |
| Firmicutes_I | Paenibacillaceae | 595 | 63 | 64 | 18 | 65 | 15 | 6 | 2 | 0 | 11 | 3 |
| Firmicutes | Staphylococcaceae | 168 | 39 | 68 | 3 | 26 | 9 | 14 | 2 | 0 | 0 | 2 |
| Proteobacteria | Moraxellaceae | 201 | 8 | 104 | 18 | 32 | 0 | 1 | 23 | 27 | 0 | 0 |
| Bacteroidota | Tannerellaceae | 542 | 121 | 34 | 3 | 8 | 23 | 1 | 0 | 0 | 0 | 1 |
| Proteobacteria | Pseudomonadaceae | 369 | 22 | 172 | 9 | 77 | 1 | 2 | 18 | 62 | 1 | 3 |
| Firmicutes | Amphibacillaceae | 203 | 38 | 70 | 0 | 31 | 1 | 3 | 1 | 0 | 8 | 0 |
| Firmicutes | Bacillaceae | 78 | 6 | 20 | 0 | 6 | 4 | 3 | 4 | 0 | 6 | 5 |
| Firmicutes | Bacillaceae_G | 38 | 3 | 11 | 4 | 20 | 0 | 6 | 4 | 0 | 2 | 4 |
| Proteobacteria | Xanthobacteraceae | 230 | 10 | 46 | 4 | 37 | 1 | 3 | 10 | 66 | 9 | 1 |

While cytochrome P450s are one of the most important and well-studied xenobiotic detoxification systems in humans, we saw relatively few homologs in gut microbes. Their homologs were also mainly restricted to facultative anaerobes, which makes sense given the oxygen-dependent mechanisms of these proteins. Furthermore, the species with the most cytochrome P450 homologs were in low-prevalence families like the *Paenibacillaceae* (**Table 2**). This suggests that while cytochrome P450 homologs can be found, they may be less likely to be relevant to the adult human gut, though with the caveat that gut oxygenation can also vary across development and in disease [21]. Flavin-dependent monooxygenases and UDP-glucuronosyltransferases had similarly sparse distributions in mostly lower-abundance gut microbes.

Two enzyme types had intermediate distributions. First, glutathione S-transferase homologs were much more commonly detected than cytochrome P450s, but were almost exclusively found in Proteobacterial facultative anaerobes, like the *Enterobacteriaceae* and *Burkholderiaceae*. However, while these are typically low-abundance, *Enterobacteriaceae* are prevalent, and Proteobacteria can rise to high levels in certain individuals and situations, making them major contributors to functional variability [22]. This suggests that gut microbial GST activity might also be especially variable across individuals. Second, arylamine acetylases were most observed in facultative anaerobes (*Enterobacteriaceae* and *Staphylococcaceae*), yet were also detected in certain *Lachnospiraceae*, the most prevalent gut microbial family worldwide [20]. Substrates for these genes include the anti-hypertensive vasodilator hydralazine [23] and the anti-tubercular isoniazid [24]. Interestingly, it has been previously shown that isoniazid is also metabolized by strains of *M. tuberculosis* by an arylamine acetylase homologous to human NAT2 [25]. Finally, arylamine acetylases are also responsible for both increasing and decreasing the carcinogenicity of certain environmental pollutants, suggesting that gut microbes could also modulate these risks [26].

In contrast, we detected thousands of short-chain and aldo-keto reductases in common gut microbes, like *Lachnospiraceae, Enterobacteriaceae*, and *Bacteroidaceae* (**Table 3**). In humans, both classes of enzymes act on a wide range of substrates; notably, certain members can participate in the reduction of steroid-like and polycyclic molecules, including bile acid intermediates [27,28]. Gut microbes are known for their ability to transform primary bile acids into secondary bile acids, and this metabolism has well-studied consequences for immune and metabolic signaling in the host [29]. Additionally, the *Lachnospiraceae* member *Clostridium bolteae* was recently found to directly metabolize the steroids nabumetone, hydrocortisone, and tacrolimus via the gene DesE [30]. We detected that the UHGP-90 protein with the best hit to DesE (GUT_GENOME228173_01934) appeared to be a full-length homolog of the human protein PECR, a trans-2-enoyl-CoA reductase that is a member of the SDR family. The prominence of reductases in the most common gut microbes aligns with previous observations that reduction reactions are especially common ways for gut microbes to transform xenobiotics, potentially because of the need for alternative electron acceptors in the absence of molecular oxygen [31,32].

Homologs of two other redox-active enzyme classes, aldehyde dehydrogenases and quinone oxidoreductases, were also observed frequently in *Lachnospiraceae*, but the highest number of distinct bacterial homologs were found in facultative anaerobes like *Enterobacteriaceae, Lactobacillaceae*, or *Burkholderiaceae*. Enzymes in these families, of course, play roles in both central and xenobiotic metabolism, complicating their interpretation. For example, aldehyde dehydrogenase oxidizes acetaldehyde to acetate (or the reverse, in microbial ethanol production). However, aldehyde dehydrogenase enzymes can have a variety of other substrates (e.g. lactaldehyde [33]) and are also involved in the detoxification of drugs like cyclophosphamide [34].

Finally, homologs of type B carboxylesterases and the “GDXG” group of lipases (which include hormone-sensitive lipases, arylacetamide deacetylases, and neutral cholesterol esterases) were also found frequently, especially in *Lachnospiraceae* and *Bacteroidaceae*. In humans, in addition to deactivating drugs like flutamide [35] and indiplon [36], these hydrolases bioactivate a large number of prodrugs, including enalapril [37] and irinotecan [38]. Overall, this analysis shows that while several systems used by humans to detoxify pharmaceutical, dietary, and environmental compounds do have at least some analog in the gut microbiome, certain enzyme families are much better represented in the most prevalent gut microbes. Specifically, these include redox-active enzymes, especially short-chain and aldo-keto reductases, and hydrolases, including lipases and carboxylesterases.

### Identifying split and full-length homologs of specific drug-metabolizing genes

In many cases, we know the specific substrates on which drug-metabolizing enzymes act. We therefore used the database PharmGKB [39] to identify cases where human proteins with gut homologs were known to be involved in the metabolism of either a pharmaceutical drug or one of its downstream metabolites.

Out of 154 proteins in PharmGKB with reviewed entries in UniProt, we found that a large majority (126/154, 82%) had at least one full-length or split homolog. 97% of these (122/126) had more full-length than split gut homologs; this set of proteins metabolized 215 drugs in total (**Supplementary Table 3**). Consistent with the sparse distribution we observed above, cytochromes were the most under-represented category in this list, with only eight genes found to have gut microbial homologs compared to 22 in PharmGKB as a whole. In contrast, 12 out of 13 drug-metabolizing UDP-glucuronosyl-transferases and all nine aldo-keto reductases were found to have full-length gut homologs.

Interestingly, despite the strong enrichment we observed for xenobiotic metabolism among proteins with full-length homologs, <50% of these proteins (56/122) fell into one of the ten classes listed above. While many different types of enzymes were represented among the remainder, a plurality of 36 were annotated in GO as metabolizing nucleobase-containing compounds. This is consistent with the observation that this process was enriched among both full-length and split homologs. Furthermore, nucleobase-containing analogs are some of the most common human chemotherapeutic, immunomodulatory, and antiviral drugs, and their metabolism is also well-studied, as variants that affect their metabolism have large consequences for health. Finally, of the remaining 30 genes, 20 were annotated in GO as oxidoreductases, further underscoring the importance of redox-active genes in gut microbial metabolism.

When we instead kept only cases with more split than full-length homologs, we found four genes involved in the metabolism of 12 drugs (**Table 4**). Again, three of these genes were involved in nucleotide metabolism, and many of these drugs were antimetabolite chemotherapeutics such as thioguanine, doxorubicin, and mercaptopurine. We noted that two out of four genes metabolized 5-fluorouracil (dihydropyrimidine dehydrogenase, or DPYD; and uridine monophosphate synthase, or UMPS). These results indicate that both full-length and split gut homologs may play roles in the microbial transformation of nucleoside analogs.

**Table 4:** Table showing all identified drug-metabolizing enzymes with more partial (nPart) than full-length (nFull) gut microbial homologs, together with the drug(s) they metabolize (From).

| Enzyme | Description | nPart | nFull | From |
| --- | --- | --- | --- | --- |
| AOX1 | Aldehyde oxidase | 46 | 3 | crizotinib, allopurinol, ziprasidone, vortioxetine, aciclovir, pyrazinamide, capmatinib, nicotine iminium ion, thioguanine |
| XDH | Xanthine dehydrogenase / oxidase | 38 | 0 | doxorubicin, 1-methylxanthine, theophylline, allopurinol, pyrazinoic acid, pyrazinamide, mercaptopurine, thioxanthine |
| DPYD | Dihydropyrimidine dehydrogenase | 26 | 24 | fluorouracil |
| UMPS | Uridine 5'-monophosphate synthase | 22 | 0 | fluorouracil |

The genes dihydropyrimidine dehydrogenase (DPYD) and xanthine dehydrogenase (XDH) had among the most split homologs. In the case of XDH, human gut microbes have been shown to catabolize purines such as uric acid, an endogenous substrate of human XDH, and to alter purine levels *in vivo* in a mouse model; knockout experiments suggest that both phenomena require an operon bearing an XDH homolog [40]. This potential conservation of function supports a potential role for XDH homologs in the metabolism of purine analog drugs, such as azathioprine (AZA), a chemotherapeutic and immunosuppressive drug that is given orally. Indeed, it has recently been shown in an *in vivo* preclinical model that *Blautia wexlerae* reduces the therapeutic effect of AZA by metabolizing its active metabolite, 6-mercaptopurine (6MP) into the inactive form, 6-TX; furthermore, this metabolism can be interrupted with the XDH inhibitor allopurinol [41].

Recent publications also support that both the human DPYD protein and its bacterial counterparts, PreT and PreA (encoded by the *preTA* operon) can inactivate the chemotherapeutic drug 5-fluorouracil (5-FU) *in vivo*. While 5-FU itself is not given orally, the orally-available prodrug, capecitabine, can also be activated to 5-FU by host liver enzymes as well as select gut bacterial strains [2]. Indeed, in a mouse model of colorectal cancer treatment with 5-FU, mice monocolonized with a *preTA* knockout strain of *E. coli* had better survival than those monocolonized with a *preTA* overexpression strain [2].

## Discussion

We conducted a systematic survey of homology between the human and gut microbial proteomes. This analysis included both full-length and “split” homologs. We found that around one in ten human proteins (2.6K) had at least some homolog in the gut microbial proteome, and that 23 human proteins had primarily split homologs. While our focus was on drug and xenobiotic metabolism, such a map of host-microbial homology may also be helpful to microbiome researchers more broadly. With this in mind, our code and results are available as publicly available resources (see **Data Availability**).

Among human proteins with full-length homologs, xenobiotic metabolism was the most enriched process, and many different xenobiotic enzyme classes were found in gut genomes. However, the most predominant systems in humans (cytochrome P450s, glutathione-S-transferases) were relatively rare in the gut microbiome. Instead, reductases and hydrolases were the most common, especially among the most prevalent microorganisms. This is consistent with previous observations about types of drug metabolism engaged in by the gut microbiome [31,32], and builds on these observations by enumerating specific classes of enzymes that are likely to contribute. Proteins involved in central metabolism, especially of nucleoside-containing compounds, were also commonly found as both full-length and split homologs in the gut microbiome. Finally, in two cases (6MP and 5-FU), the gut microbial split homologs we identified have been shown to metabolize pharmaceutical nucleoside analogs in mouse models [2,41].

One limitation of this work is that we have only considered the gut microbiome. This community was our focus because microbial biomass is highest in the gut [42], because orally ingested drugs are absorbed in the intestine [43], and because the gut is closely connected to the liver, the primary site of drug metabolism in humans [44]. However, split orthologs in skin microbes may also be relevant for topically applied drugs, and similarly for oral microbes and drugs delivered as rinses.

A technical limitation of this work is that sequencing and annotation errors can give rise to *in silico,* artifactual gene “fusions” or “fissions.” We believe that the way that UHGP-90 protein clusters were constructed would favor such “fusions.” UHGP-90 protein clusters were constructed using MMSeqs2’s “linclust” algorithm [45] in target-coverage mode, meaning that the representative sequence for a cluster must cover 80% of each member sequence, but not necessarily vice versa. This has the advantage that protein fragments or artifactual “fissions” would seldom be chosen as representative sequences, but also means artifactual “fusions” would be chosen more often. Since we use the representative sequences in this pipeline, this effect would therefore bias us away from detecting split homologs.

Of course, it is important to emphasize that homologs may differ in substrate specificity. This is especially true over long evolutionary distances (e.g., between humans and microorganisms) and for enzymes whose substrate specificity is broad (e.g. many xenobiotic metabolism genes). Follow-up experiments would therefore be necessary to establish whether specific substrates are shared between host and microbial homologs. Advances in computational structural biology, such as improvements to high-throughput ligand docking tools [46–48], may also help prioritize homologs that could contribute to parallel drug metabolism between host and microbiome. We speculate that interactions between central metabolic enzymes and substrate analogs, such as the chemotherapeutics 6MP and 5-FU, may be especially likely to translate: these enzymes are more evolutionarily constrained than broad-spectrum xenobiotic enzymes [9], and the corresponding drugs bind in the active site, which is typically highly conserved.

While we have focused on proteins that metabolize drugs, the protein targets of drugs could also be conserved, potentially causing off-target effects on the microbiome. Such unintended effects of pharmaceuticals on gut microorganisms are not rare: one study of more than 1,000 marketed drugs found that nearly a quarter inhibited the growth of at least one of 40 representative gut isolates [49]. As above, we would expect host-microbiome homology to be especially relevant when considering proteins targeted by substrate analogs. Indeed, a study of the chemotherapeutic 5-FU showed that it had large effects on gut microbial growth [2], and antimetabolites as a class were also enriched for antimicrobial effects in the study above [49].

A final unresolved question is the evolutionary history of these homologs. For example, which of these homologs are descendants of ancestral sequences present in the last universal ancestor, and which might be better explained by horizontal gene transfer? Transfers from bacteria into early eukaryotes [12], especially from the ancestors of modern organelles [50], as well as transfers from modern eukaryotes into bacteria [51], are two mechanisms that could lead to both full-length and split homology. While ancient events are intrinsically difficult to resolve, phylogenetic methods, combined with the current explosion in microbial genome sequencing, may enable us to distinguish between these possibilities.

## Methods

### Identification of split and full-length homologs

To identify homologs, we performed a BLASTP search of all 13.9M proteins in the UHGP-90 database against all 20.6K human proteins downloaded from UniProt (2023/9/13). This yielded 8.5M potential matches. We then performed the following filtering steps:

1. <u>Best human hit:</u> for each UHGP-90 protein, retain only the human protein with the highest bitscore;
2. <u>Microbe coverage:</u> retain only alignments covering at least 67% of the prokaryotic UHGP-90 protein sequence; additionally, retain only UHGP-90 sequences that are at least 80 amino acids long.

The pipeline diverged after this point for full-length and split homologs. For full-length homologs, we were interested in individual microbial proteins where an alignment covered most of the human protein, and where this alignment was unlikely to be due to contamination in the microbial genomes. We therefore performed the following filtering steps:

F3. <u>Human coverage:</u> each alignment must cover at least 70% of the human sequence;

F4. <u>Contamination:</u> filter out any alignments whose amino acid percent ID was more than three standard deviations above the mean (mean: 30% ID; cutoff: 51.8% ID).

For split homologs, we sought to determine which UHGP-90 families were encoded by neighboring features in the same genome, and where this was unlikely to be the result of contamination, as above. Because multiple genomes could encode the same UHGP-90 family, we had to first expand our results, then filter, as follows:

S3. <u>Same genome:</u> first, determine which individual UHGG genomes encoded multiple UHGP-90 families aligning to each individual human protein. Then, retain only those alignments, repeated for each genome that encoded the UHGP-90 protein;

S4. <u>Joint human coverage:</u> for each UHGG genome and for each human protein, determine whether the alignments between the human protein and the UHGP-90 proteins from that genome could jointly, but not individually, cover at least 70% of the human sequence;

S5. <u>Feature distance:</u> for each UHGG genome and for each human protein, compute the minimum distance (in feature numbers) between each UHGP-90 protein aligning to that human protein. Retain only sets of alignments with at least two different UHGP-90 proteins that are three or fewer features apart, and that are additionally all on the same strand and contig. Then, repeat the human coverage step to ensure that the remaining alignments still jointly cover ≥70% of the human protein, as some have been removed;

S6. <u>Contamination:</u> filter out any alignments whose amino acid percent ID was more than three standard deviations above the mean observed for all full-length homologs, as above (mean: 30% ID; cutoff: 51.8% ID); repeat the human coverage step again and report results.

Filtering and analysis steps were carried out in a Snakemake pipeline, using Pandas [52], Polars [53], and R with Tidyverse [54,55].

### Enrichment analysis

Gene Ontology (GO) annotations [17] from UniProt [15] were used to determine subcellular localization (“cellular component”) and function (“biological process”) for human proteins.

Proteins whose cellular component annotations matched the regular expression “[Mm]itochondr” were retained as mitochondrially-localized. Enrichment analysis was carried out using TopGO [56] using Fisher’s exact test on GO biological process terms, with the resulting *p*-values corrected for multiple testing using the Benjamini-Hochberg method [57].

### Analyzing subclasses of xenobiotic enzyme families

Xenobiotic enzyme families were defined using protein family annotations in UniProt, using regular expression matches for the following:

- <u>Aldo-keto reductases</u> (“akr”): “Aldo/keto reductase family”;
- <u>UDP-glucuronosyltransferases</u> (“udp”): “UDP-glycosyltransferase family” (note: all human members of this family except cerebroside synthase were annotated as UDP-glucuronosyltransferases);
- <u>Glutathione S-transferases</u> (“gst”): “GST superfamily” (note: this included the alpha, zeta, sigma, pi, mu, theta, omega, and kappa families);
- <u>Arylamine N-acetyltransferases</u> (“aryl”): “Arylamine N-acetyltransferase family”;
- <u>GDXG-like hydrolases</u> (“gdxg”): “’GDXG’ lipolytic enzyme family”;
- <u>Cytochrome P450s</u> (“cyto”): “Cytochrome P450 family”;
- <u>Type B carboxylesterases</u> (“ester”): “Type-B carboxylesterase/lipase family”;
- <u>Flavin monooxygenases</u> (“flavin”): “Flavin monoamine oxidase family|FMO family” (note: this included the FIG1 subfamily);
- <u>Short-chain reductases</u> (“sdr”): “Short-chain dehydrogenases/reductases (SDR)”;
- <u>Quinone oxidoreductases</u> (“quin”): “Quinone oxidoreductase subfamily”.

Full-length homologs were partitioned into one of the above classes. Next, for each class, the phylogenetic diversity of species containing at least one full-length homolog was calculated using Faith’s PD [58]. The phylogeny used was the maximum-likelihood tree of the 4,616 species in UHGG [14] generated via IQ-TREE [59], which we midpoint-rooted using APE [60]. Faith’s PD was calculated using Picante [61]. Xenobiotic classes were visualized in descending order of PD. The number of unique species with at least one full-length homolog is given in **Table 2**, while the total number of unique UHGP-90 IDs per family is given in **Table 3**. Results were visualized using the R package ggtree [62].

### Identification of gut homologs of drug-metabolizing enzymes

Pathway, gene, relationship, and chemical annotations were downloaded from PharmGKB (2024/10/12) [63]. HUGO Gene Nomenclature Committee (HGNC) identifiers [64] in PharmGKB were mapped, using data downloaded from HGNC (2024/10/01), to UniProt IDs. Chemicals of interest in PharmGKB were defined as having the chemical classes “Drug”, “Drug Class”, “Prodrug”, or “Metabolite” (this refers to drug metabolites, not endogenous substrates or “Biological Intermediates”, which were excluded). This was necessary because certain endogenous human metabolites were annotated in pathways, but only peripherally related to drug metabolism, e.g., homocysteine in the methotrexate metabolism pathway (present because methotrexate targets folate biosynthesis, which in turn is linked to homocysteine via the S-adenosyl-methionine cycle). Reactions from all PharmGKB pathways were then filtered such that the reactant and products were different, the reaction type was not “Transport”, the “Controller” (typically an enzyme or regulator) was known, and either the reactant or product (or both) was a chemical of interest as defined above.

### Identification of other xenobiotic enzyme types in PharmGKB

In addition to the xenobiotic enzyme classes we defined above, we also considered two other classes of enzymes:

- <u>Nucleobase metabolism genes</u>: genes annotated to the GO term “nucleobase-containing compound metabolic process (GO: 0006139)”, or any term below it, but excluding genes annotated to the following GO terms or any terms below them:

- “Acyl-CoA metabolic process (GO:0006637)”;
- “Coenzyme A metabolic process (GO:0015936)”;
- “FMN metabolic process (GO:0046444)”;
- “FAD metabolic process (GO:0046443)”;
- “Pyridine nucleotide metabolic process (GO:0019362)”.
- <u>Oxidoreductases</u>: genes annotated to the GO term “oxidoreductase activity (GO:0016491)”.

### Identification of DesE homologs

In the study showing DesE from *Clostridium bolteae* metabolized the ketone group of pharmaceutical steroids [30], its GenBank [65] protein accession was given as EDP16280.1. To determine whether this protein was represented in our full-length homologs, we first retrieved this accession and used it to perform an MMSeqs2 [45] search in “easy-search” mode against all UHGP-90 protein sequences. The best-hit protein (GUT_GENOME228173_01934, 97.7% identity) was then used to filter our list of full-length homologs. GUT_GENOME228173_01934 was found to be a full-length homolog of the human gene PECR (UniProt ID Q9BY49), and was distributed in six Lachnospiraceae species; these included *Clostridium_M bolteae* where it was discovered and two other species in the genus *Clostridium_M*.

## Data availability

The UHGG and UHGP datasets [14] can be obtained from EMBL-EBI [66] at https://ftp.ebi.ac.uk/pub/databases/metagenomics/mgnify_genomes/human-gut/v1.0/. The human reference proteome was downloaded from UniProt [15] and is available at https://ftp.uniprot.org/pub/databases/uniprot/previous_releases/release-2023_03/. HGNC IDs [64] can be obtained at https://storage.googleapis.com/public-download-files/hgnc/archive/archive/monthly/tsv/hgnc_complete_set_2024-10-01.txt.

Code for our pipeline and analysis are available at https://github.com/pbradleylab/split_homology. The processed output of the pipeline (described starting on p. 13) is available via Zenodo at https://zenodo.org/uploads/14037045 (DOI: 10.5281/zenodo.14037045).

## Acknowledgements

The authors wish to thank Abigail Lind for helpful feedback. Funding was provided by The Ohio State University (P.H.B.) and the National Institutes of Health: R35GM151155 (P.H.B.); R01CA255116, R01HL122593 (P.J.T). P.J.T is a Chan Zuckerberg Biohub-San Francisco Investigator.

## Supplementary Tables and Figures

**Supplementary Figure 1:**
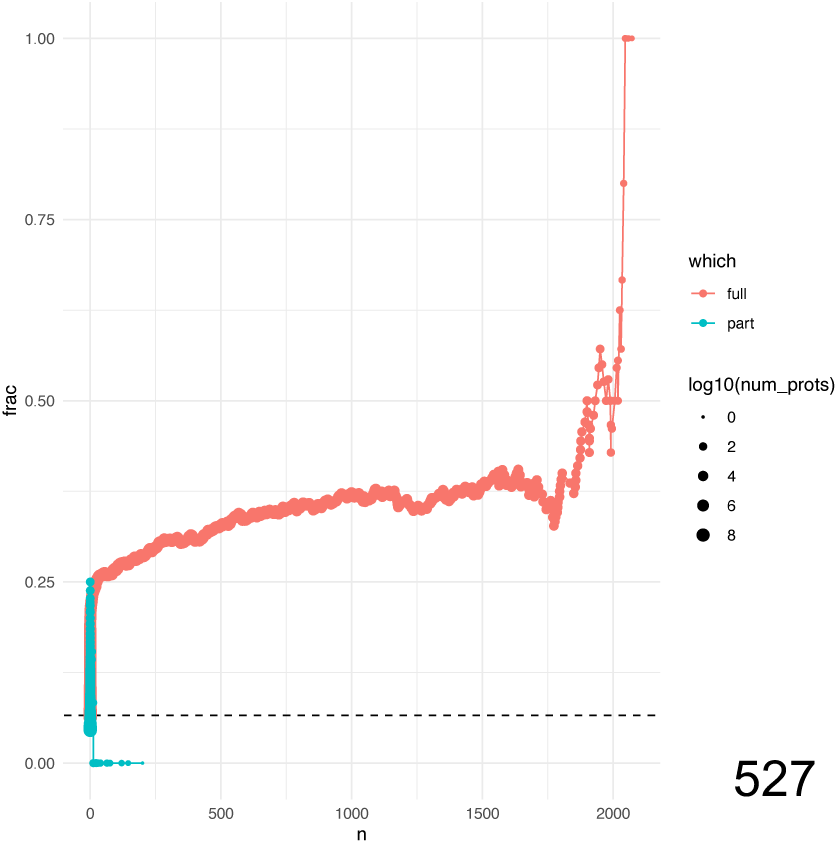
Trend in mitochondrial localization for full-length (orange) and split (teal) homologs, as a function of their distribution across gut species. Each dot represents all microbial homologs present in at least a certain number of gut microbial species (x-axis). The size of the dot corresponds to the total number of such microbial homologs. The y-axis shows what fraction of the human homologs are annotated as localizing to the mitochondrion. The overall rate for all proteins is shown by the dashed line.

**Supplementary Table 1:** GO term enrichment (biological process) for human proteins with more full-length than split homologs. Terms with adjusted p-values below 0.05 are shown.

**Supplementary Table 2:** GO term enrichment (biological process) for human proteins with more split than full-length homologs. Terms with adjusted p-values below 0.05 are shown.

**Supplementary Table 3:** Drugs metabolized by human proteins with mostly full-length homologs in the gut microbiome.

